# Regolith as a Refuge: Differential Survival of Bacteriophage Qβ in Mars-Analog Environments

**DOI:** 10.64898/2025.12.02.691594

**Authors:** Miguel Arribas Tiemblo, Alicia Rodríguez-Moreno, Felipe Gómez, Ester Lázaro

## Abstract

Viruses are among the simplest biological entities capable of replication. Their robustness and adaptability make them relevant not only to terrestrial ecosystems but also to astrobiological exploration. As durable entities, they may persist in environments far harsher than those tolerable to cellular life and are likely candidates for forward contamination. To assess their relevance in planetary protection and as potential biomarkers, we investigated the preservation of bacteriophage Qβ, an RNA virus, in Mars-analog environments using two commercial Martian regolith simulants: MGS-1 and MMS-2. MMS-2, enriched in iron oxides, exhibited higher oxidative potential than MGS-1, which is mainly composed of basaltic material. Viral survival was assessed across variables including time, temperature, concentration, and particle size. Our results show that inactivation in MGS-1 was primarily driven by adsorption, while in MMS-2 it was dominated by chemical oxidation. MGS-1 provided a more protective matrix, mitigating freeze-induced damage and shielding desiccated viral particles from UV-B and UV-C radiation. These findings highlight the importance of mineral composition in modulating viral persistence and suggest that regolith may act as both a barrier and a refuge. Understanding virus–mineral interactions is essential for assessing biosignature preservation, planetary protection, and the potential roles of viruses in the evolution and survival of life beyond Earth.

## Introduction

The effects of extreme environmental conditions on microorganisms, −including high salt concentration, limited water availability, high radiation levels, extreme temperatures, and atypical gravity− have been extensively studied, both in natural settings and under controlled laboratory conditions (Fukunaga, 2020; Horneck, 1981; Horneck et al., 2010; Moeller et al., 2010). Some of these extreme conditions serve as analogues for planetary scenarios (Rettberg et al., 2004). In particular, the survival of microorganisms in simulated Martian habitats has been the focus of several recent studies (Beblo-Vranesevic et al., 2022; Horne et al., 2022).

The global mean annual surface pressure on Mars is approximately 6 hPa −significantly lower than the 1013 hPa at sea level on Earth (Haberle, 2015). In addition, the Martian surface is exposed to intense and biologically harmful radiation (Zaccaria et al., 2024). Depending on latitude, the surface receives between 7–10 W/m² of UVA, 2–5 W/m² of UVB, and 0.5–1.5 W/m² of UVC radiation at noon (Rodriguez-Manfredi et al., 2021). The total dose rate of ionizing radiation reaches approximately 233 ± 12 µGy/day (Matthiä et al., 2017; Zeitlin et al., 2019), which is about ten times higher than that on Earth. The intensity of this radiation can also lead to secondary effects through the generation of reactive oxygen species (ROS) and Fenton reactions (Bagnato et al., 2021; Edgar et al., 2022). Due to the high cost of exploratory missions and the lack of direct Martian soil samples on Earth −aside from meteorites− researchers have turned to terrestrial analog environments to simulate Martian conditions (Shen et al., 2022). Examples include the Atacama Desert in Chile (27° 22′ 00″ S, 70° 19′ 56″ W), known for its extreme aridity and intense UV radiation, and the Río Tinto region in Huelva, Spain (37° 42′ 11″ N, 6° 33′ 10″ W), which shares mineralogical similarities with Martian terrain (Amils et al., 2023; Gómez et al., 2024).

In addition, commercial regolith simulants analogous to Martian soil −such as Mars Global Simulant (MGS-1) (Cannon et al., 2019) and Mojave Mars Simulants (MMS-1 and MMS-2) (Peters et al., 2008)− have been developed to replicate the physical and chemical properties of Martian surface materials. These simulants are typically derived from basaltic terrains and are designed to mimic the mineralogical composition and particle size distribution of Martian regolith as closely as possible. MGS-1 is modeled after the Rocknest aeolian deposit at Gale Crater, incorporating data from both landed and orbital missions. It is currently the most thoroughly characterized Martian regolith simulant (Karl et al., 2022). In contrast, MMS-1 and MMS-2 are less commonly used alternatives (Clark et al., 2020). MMS-1 was originally developed by the Jet Propulsion Laboratory (JPL, Pasadena, California, USA) from a basaltic flow in the Mojave Desert (Caporale et al., 2020). According to the supplier’s specifications, MMS-1 is chemically analogous to the original JPL simulant, while MMS-2 is a modified version enriched with iron and magnesium oxides, silica sand, and gypsum to better reflect the soil composition measured by the Opportunity rover, aiming to represent the average Martian surface (Costa et al., 2024).

The preservation of biomarkers within these regolith simulants has been widely studied under Martian simulated surface conditions. Amino acids, for instance, have variable half-lives in the Martian surface that heavily depend on the nature of their side chain (Laurent et al., 2019; dos Santos et al., 2016). Their half-lives increase by over four orders of magnitudes when protected from radiation by the regolith (ten Kate et al., 2006, 2005). The detection and characterization of these biosignatures can be, however, hindered by the regolith itself, as the presence of iron oxides like magnetite, adsorptive processes and substrate-induced oxidation can all negatively affect the detection and recovery of organics from Martian regolith simulants (Tiemblo et al., 2025). Other biomarkers, like cyanobacterial carotenoids, lipids and microorganisms themselves are also better preserved when within these simulants, although detection within them is systematically complex (Baqué et al., 2016; Billi et al., 2022, 2019; Fagliarone et al., 2020; Rzymski et al., 2022). The survivability and ease of detection of diverse microorganisms have also been studied under Mars Surface conditions. Facultative anaerobes from analogue sites have been employed to simulate potential subsurface habitable niches (Beblo-Vranesevic et al., 2022). Cyanobacteria under perchlorate, atmospheric and UV stress can not only survive but actively replicate in the appropriate conditions (Arribas Tiemblo et al., 2025; Rzymski et al., 2022). More recently, human-associated pathogens have also been tested under Mars-analogue conditions (Zaccaria et al., 2024), revealing that certain terrestrial microorganisms can survive in simulated Martian settings.

In contrast, our understanding of how such extreme conditions affect viral populations remains limited (Trubl et al., 2023). This gap is particularly relevant given the remarkable capacity of viruses to endure conditions that are lethal to most cellular life, which makes them uniquely robust (de la Higuera and Lázaro, 2022) and has led to speculation about their potential role as carriers of genetic material across planetary boundaries (Berliner et al., 2018; Griffin, 2013). This property, together with the extraordinary adaptability and evolutionary potential of viruses, even in conditions that life on Earth has not experienced, such as the absence of gravity (Rodríguez-Moreno et al., n.d.), make them compelling systems for studying the persistence of simple biological systems under conditions typical of other planetary environments.

Viruses are the most numerous biological entities on Earth and are present in virtually all ecosystems, including extreme environments considered analogues of other planetary settings (Berliner et al., 2018). Consequently, they may also represent a potential source of forward contamination during crewed missions to Mars. As the prospect of long-duration and increasingly distant space exploration becomes more tangible, it is essential to address the microbial risks associated with such missions (Trubl et al., 2023), not only in terms of astronaut health and safety, but also with regard to the integrity of the planetary environments to be explored. The development of comprehensive bioburden assessment protocols that explicitly include viruses is therefore a critical step toward effective planetary protection.

Among viruses, bacteriophages (or phages) are especially abundant, outnumbering their bacterial hosts in most ecosystems. Their simplicity and lack of biosafety restrictions make them ideal candidates for experimental studies on viral survival under extreme conditions, including those that simulate environments of astrobiological interest. In this work, we use the bacteriophage Qβ, a positive-sense single-stranded RNA virus that specifically infects *Escherichia coli* strains expressing conjugative pili (F⁺ strains). First isolated in 1961 from human feces (Arisaka, 2024), Qβ can be considered a potential component of the gut microbiome (Bastin et al., 2020). Its host, *E. coli*, is a well-characterized microorganism commonly used in laboratory settings, further supporting the relevance of this model system. Qβ belongs to the *Fiersviridae* family (formerly *Leviviridae*) and the *Qubevirus* genus (Walker et al., 2021). The genomic RNA (gRNA) of Qβ, comprising 4,217 nucleotides, encodes the maturation protein (Mat), the major coat protein (CP), the minor capsid protein, and the β-subunit of replicase. The Mat and CP proteins are responsible for the structural integrity of the phage capsid, which has a quasi-symmetry T=3 and an average diameter of 25–30 nm (Chang et al., 2022; Cui et al., 2017; Gorzelnik et al., 2016).

The main objective of this study is to explore the interaction between bacteriophage Qβ and two Martian regolith simulants −MGS-1 and MMS-2− under a range of Mars-analog environmental conditions. By systematically assessing viral survivability across variables such as temperature, radiation, desiccation, and mineral composition, we aim to shed light on the potential fate of viral particles on the Martian surface. Our findings contribute to a deeper understanding of viral resilience in extraterrestrial environments and offer new perspectives for astrobiology, planetary protection, and the microbial risks associated with future space missions.

## Materials and methods

### Geochemical analysis

The mineralogical analysis of the Martian Regolith Simulants was performed through X-Ray Diffraction (XRD). Prior to analysis, the regolith simulants were separated into five particle size ranges (< 0.045 mm, 0.045-0.125 mm, 0.125-0-25 mm, 0.25-0-5 mm and > 0.5 mm) through highly precise sieves. Phase identification was performed using DIFFRACT.EVA software in conjunction with the PDF-2 mineral database. Semiquantitative phase analysis was conducted using the Reference Intensity Ratio (RIR) method in a D8 Advance ECO X-Ray Diffractometer (Bruker), in which the *I/Ic* parameter is defined as the ratio between the intensity of the most prominent diffraction peak of the phase of interest and that of the most intense peak of corundum. RIR values for a wide range of crystalline phases are available in the reference database employed. In cases where the identified phase lacked a tabulated *I/Ic* value, a default value of 1 was assigned. The calculation assumes that all crystalline phases present in the sample have been correctly identified; consequently, the software normalizes the results such that the sum of all phase concentrations equals 100% (∑ cᵢ = 100%).

### Culturing of viruses

The plasmid pBRT7Qβ, which contains a cDNA of bacteriophage Qβ cloned in the plasmid pBR322 (Barrera et al., 1993; Taniguchi et al., 1978), was used to transform *Escherichia coli* DH5-α. This strain allows for viral expression but cannot be infected due to the absence of the F pilus. The supernatant of an overnight culture obtained from a transformed colony was used to infect *E. coli*, strain Hfr (Hayes, 1953), in semisolid agar. A lysis plaque was randomly selected, and the resulting viral progeny was amplified by infecting an exponential-phase *E. coli* Hfr culture with 10⁶ plaque-forming units (PFU) under standard laboratory conditions (37 °C, 250 rpm for 2 h in NB medium: 8 g/l Nutrient Broth from Merck and 5 g/l NaCl) using a New Brunswick Scientific Innova 42 Incubator Shaker (Eppendorf, Enfield, CT, USA). After incubation, the culture was treated with chloroform (1:20 v/v, 28 °C, 15 min, 850 rpm in a thermomixer) and centrifuged at 13,000 rpm. The resulting supernatant, containing viral particles, was used as the working viral stock throughout this study. For each experimental assay, fresh dilutions of this stock were prepared in phage buffer (25 mM Tris-HCl, pH 7.5; 5 mM MgCl₂; 1 g/l gelatin) to an approximate concentration of 10^8^ PFU/ml.

### Standard procedures for plaque assays

Bacteria used for plaque assays (*E. coli* Hfr) were obtained by inoculating a well-isolated colony into a 250 ml PYREX® Erlenmeyer flask (Schott Duran) containing 50 ml of NB medium, and incubated overnight at 37 °C in a New Brunswick Scientific Innova 42 Incubator Shaker (Eppendorf, Enfield, CT, USA) at 250 rpm. Plaque assays were performed by adding to 4.5 ml of soft agar (7 g/l agar in NB medium), 300 µl of the bacterial culture, and 100 µl of the viral supernatant diluted in phage buffer. The mixture was poured onto LB agar plates (32 g/l LB agar; Lennox L Broth Base from Invitrogen) and incubated at 37 °C overnight. Dilutions were chosen to yield between 20 and 200 lysis plaques per plate. Viral titers were calculated by counting the number of plaques and multiplying by the corresponding dilution factor. They were expressed as plaque-forming units (PFU) per ml of the phage suspension. Therefore, viral titers always refer to the number of infective particles.

### Simulation of Martian conditions through Martian Regolith Simulants

A working viral solution (approximately 10⁸ PFU/ml in phage buffer) was used for all procedures. Depending on the set-up, this solution was distributed into tubes (2 ml microcentrifuge tubes) or 96-well plates at a final volume of 200 µl. Any deviations from this standard procedure are explicitly indicated in the corresponding section.

#### Time-dependent viral inactivation

A working viral solution, prepared as previously described, was mixed with the Martian Regolith Simulants (MRSs) MGS-1 and MMS-2 at a 12.5% w/v ratio (25 mg of simulant per 200 µl of viral solution) in tubes. Mixtures were incubated at 25 °C with agitation at 800 rpm using a thermomixer (Thermomixer comfort, Eppendorf, 2 ml). At each time point, tubes were briefly centrifuged (10 s spin) to sediment the regolith simulant, and 10 µl of the supernatant were immediately used for plaque assays to quantify the remaining infective viral particles. Time points were taken at 0, 5, 10, 20, 30, and 60 minutes for MMS-2, and at 0, 5, 30, 60, and 120 minutes for MGS-1.

The loss of viral infectivity over time was modeled using a four-parameter logistic (4PL) function, which describes a sigmoid-like curve. In this model, the upper asymptote (U) was set to 1, representing the initial concentration of infective viral particles. The lower asymptote (L) corresponds to the level at which viral infectivity stabilizes after prolonged exposure (> 1 h). The slope (S) defines the steepness of the curve at the inflection point (x₅₀), which in this case represents the time required for a 50% reduction in the titers of initial viral population.

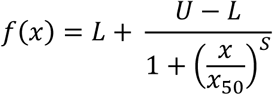

#### Concentration-dependent viral inactivation

A working viral solution in tubes was mixed with the two MRSs (MGS-1 and MMS-2) at five different regolith-to-solution ratios: 0.5%, 2.5%, 12.5%, 50%, and 100% w/v. Mixtures were vortexed briefly and incubated at 25 °C with agitation at 800 rpm for 20 minutes in the thermomixer. After incubation, samples were centrifuged at 13,000 rpm for 10 minutes, and the supernatants were immediately used for plaque assays to quantify the remaining infective viral particles.

#### Temperature-dependent viral inactivation

The working viral solution in tubes was mixed with the MRSs MGS-1 and MMS-2 at a 12.5% w/v ratio (25 mg of simulant per 200 µl of viral solution). Samples were incubated at four different temperatures −4 °C, 25 °C, 30 °C, and 37 °C− with agitation at 800 rpm in the thermomixer. For each temperature, two incubation times were tested: 20 minutes and 120 minutes. After incubation, samples were centrifuged at 13,000 rpm, and the supernatants were immediately used for plaque assays to quantify the remaining infective viral particles.

#### Assessment of different particle sizes

To assess the influence of particle size on viral inactivation, the two MRSs, MGS-1 and MMS-2, were separated into five size ranges using precision sieves: <0.045 mm, 0.045–0.150 mm, 0.150–0.250 mm, 0.250–0.500 mm, and >0.500 mm. Working viral solutions in tubes were mixed with each particle size fraction at a 12.5% w/v ratio (25 mg of simulant per 200 µl of viral solution). Mixtures were vortexed and incubated at 800 rpm in thermomixer under four conditions: 120 minutes at 4 °C, 25 °C, and 37 °C, and 20 minutes at 25 °C. After incubation, samples were centrifuged at 13,000 rpm, and the supernatants were immediately used for plaque assays to quantify the remaining infective viral particles.

#### Desiccation and UV-B/C irradiation experiments

To assess the effect of desiccation and ultraviolet (UV) irradiation on viral survival, the working viral solution was mixed with MGS-1 at a 12.5% w/v ratio (25 mg per 200 µl) into wells of a 96-well microplate. Prior desiccation, samples were incubated at 25 °C for 10 minutes with agitation at 150 rpm. Desiccation was performed at room temperature using a Concentrator plus/Vacufuge® plus (Eppendorf), operated in desiccator mode (D-AQ: desiccator–aqueous) until samples were completely dried.

Once dried, samples were subjected to UV irradiation. UV-B exposure was carried out in a BS-02 Opsytec Dr. Gröbel irradiation chamber equipped with 20 Philips TLD 15W/05 UV-B lamps. Samples were placed 28 cm below the light source and irradiated at a fluence of 7 mW/cm² until reaching a total dose of 25 J/cm², measured with a Dr. Gröbel UV-B dosimeter. UV-C irradiation was performed in the same chamber using Philips TUV 15W/G15 T8 UV-C lamps, without lamp attenuation, at a fluence of 2.59 mW/cm² and a total dose of 9 J/cm², measured with an RM12 Opsytec UV sensor.

Following irradiation, viral particles were recovered by adding 200 µl of phage buffer to each well. The recovered solution was centrifuged at 13,000 rpm for 10 minutes and the supernatants were immediately used for plaque assays to determine viral survival.

#### Resistance to Freezing and Thawing

To evaluate the resistance of viral particles to freezing and thawing (F/T) in the presence of MRSs, working solutions in tubes (in this case 1,5 ml microcentrifuge tubes) were subjected to five consecutive F/T cycles. For each assay, the working viral solution were mixed with MGS-1, MMS-2, or sea sand (functioning as inert control) at a 12.5% w/v ratio, vortexed briefly, and frozen at −20 °C. Samples were thawed at 4 °C every 24 hours, except for the final cycle, which included a 14-day freezing period. A 10 µl aliquot was taken after each thaw for plaque assay quantification of infectious virions. The remaining volume was immediately refrozen. The survival rate was calculated as the ratio between the titer obtained after the final cycle and the initial titer at the onset of the experiment.

## Statistics

Statistical analyses were performed with R (version 4.3.2). All tests were two-tailed, and statistical significance was set at *p* < 0.05. Welch’s *t*-test was used to assess differences between groups, as unequal variances were expected across conditions. In specific cases, one-way ANOVAs followed by post-hoc Dunnett’s tests were conducted. Statistical significance is indicated in the text using asterisks: (\**) p < 0.05, (**) p < 0.01, and (\*\*\**) *p* < 0.001.

All experimental points were conducted in at least triplicate. Data presented in the graphs are expressed as mean values ± standard deviations. The survival rate was defined as the proportion of viral particles that retained infectivity following exposure to environmental factors. It was calculated by dividing the number of infectious viruses after treatment (N) by that of the untreated control (N_0_) (Survival Rate = N/N_0_).

## Results

### Mineral composition of the MRSs

Bulk composition data for both MGS-1 and MMS-2 are publicly available from their respective suppliers: Exolith Lab for MGS-1 (Exolith Lab (MGS-1), 2023) and the Martian Garden for MMS-2 (The Martian Garden, 2024). Both simulants were fractionated into five particle size ranges (< 0.045, 0.045–0.150, 0.150–0.250, 0.250–0.500, and > 0.500 mm), and their mineral composition was analyzed by X-ray diffraction (XRD) (Figure 1).

**Figure 1.**
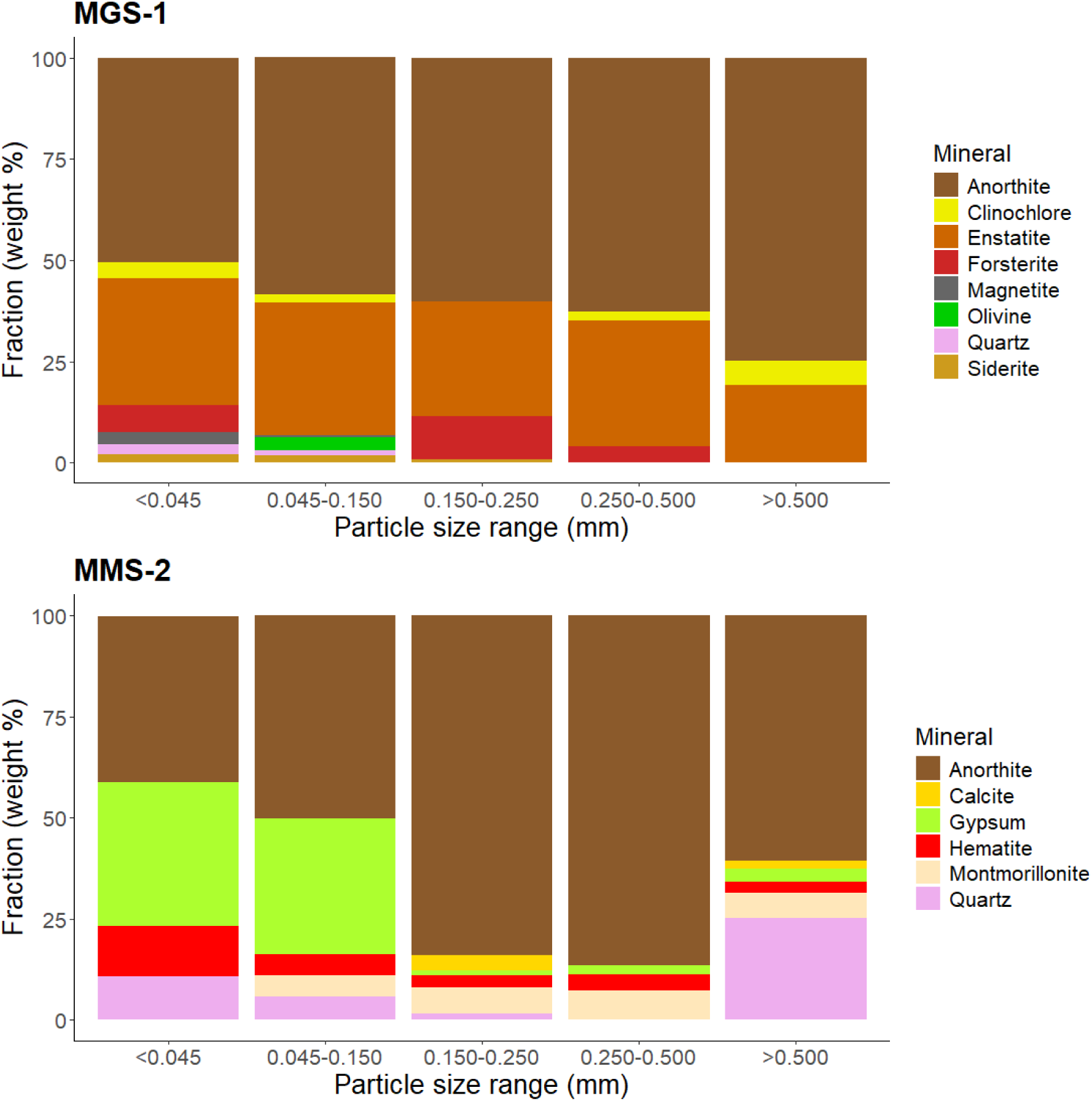
Mineral composition of the Martian Regolith Simulants MGS-1 and MMS-2, fractionated into five particle size ranges (< 0.045 mm, 0.045–0.150 mm, 0.150–0.250 mm, 0.250–0.500 mm, and > 0.500 mm). Samples were analyzed by X-ray diffraction (XRD) using a D8 Advance ECO diffractometer (Bruker). Mineral abundance is expressed as weight percentage (% w/w) of each fraction.

The main component of both simulants was anorthite-like plagioclase, a mineral commonly found in basaltic rocks (Huang et al., 2025a; Yu et al., 2016). All particle sizes of MGS-1 also contained significant amounts of enstatite, a magnesium-rich pyroxene. Minerals from the olivine group (olivine and forsterite) were detected in lower abundance, particularly in the finer fractions. Overall, MGS-1 exhibited a consistent mineralogical profile across particle sizes, with only minor variations−namely, a progressive dominance of anorthite in larger particles and a greater diversity of minerals in the finer fractions. MGS-1 was primarily composed of inert silicates, with minimal to no detectable oxides.

In contrast, the composition of MMS-2 varied markedly with particle size. Gypsum, along with anorthite, was one of the dominant mineral phases in particles smaller than 0.150 mm, but was largely absent in coarser fractions. Particles between 0.150 and 0.500 mm were predominantly composed of anorthite, while those larger than 0.500 mm contained substantial amounts of quartz, often appearing as visible chunks (> 1 mm). Hematite, an iron oxide, was abundant in the finest fraction (< 0.045 mm), but was present across all size ranges at concentrations exceeding 3% w/w. Montmorillonite, a smectite clay, was also consistently detected in all fractions > 0.045 mm.

### Phague - MSRs interaction

Viral solutions were exposed to the two Martian Regolith Simulants, MGS-1 and MMS-2, to evaluate the effects of parameters relevant to the Martian surface environment.

#### Variations in exposure time

The loss of viral infectivity (defined as survival rate) upon mixing the virus solution with both MRSs followed a reverse sigmoid function, modeled parametrically (Figure 2A). Although viruses were incubated alongside MGS-1 for up to 120 min, data is only shown up to 90 min, as all relevant changes happen in the first 30 min or so. Exposure to MMS-2 resulted in near-complete viral inactivation (>99%), significantly higher than that observed with MGS-1 (∼75%) (Figure 2B). However, 50% inactivation occurred more rapidly in MGS-1 (5 min) than in MMS-2 (13 min) (Figure 2C). The slopes of both sigmoid curves were comparable, and they intersected at 20 minutes of exposure, corresponding to ∼75% inactivation. Prior to this point, MGS-1 induced greater viral loss; beyond it, MMS-2 was more harmful. Based on these observations, a 20-minute incubation period was selected for subsequent experiments, as it allowed for consistent and comparable assessments of both simulants.

**Figure 2.**
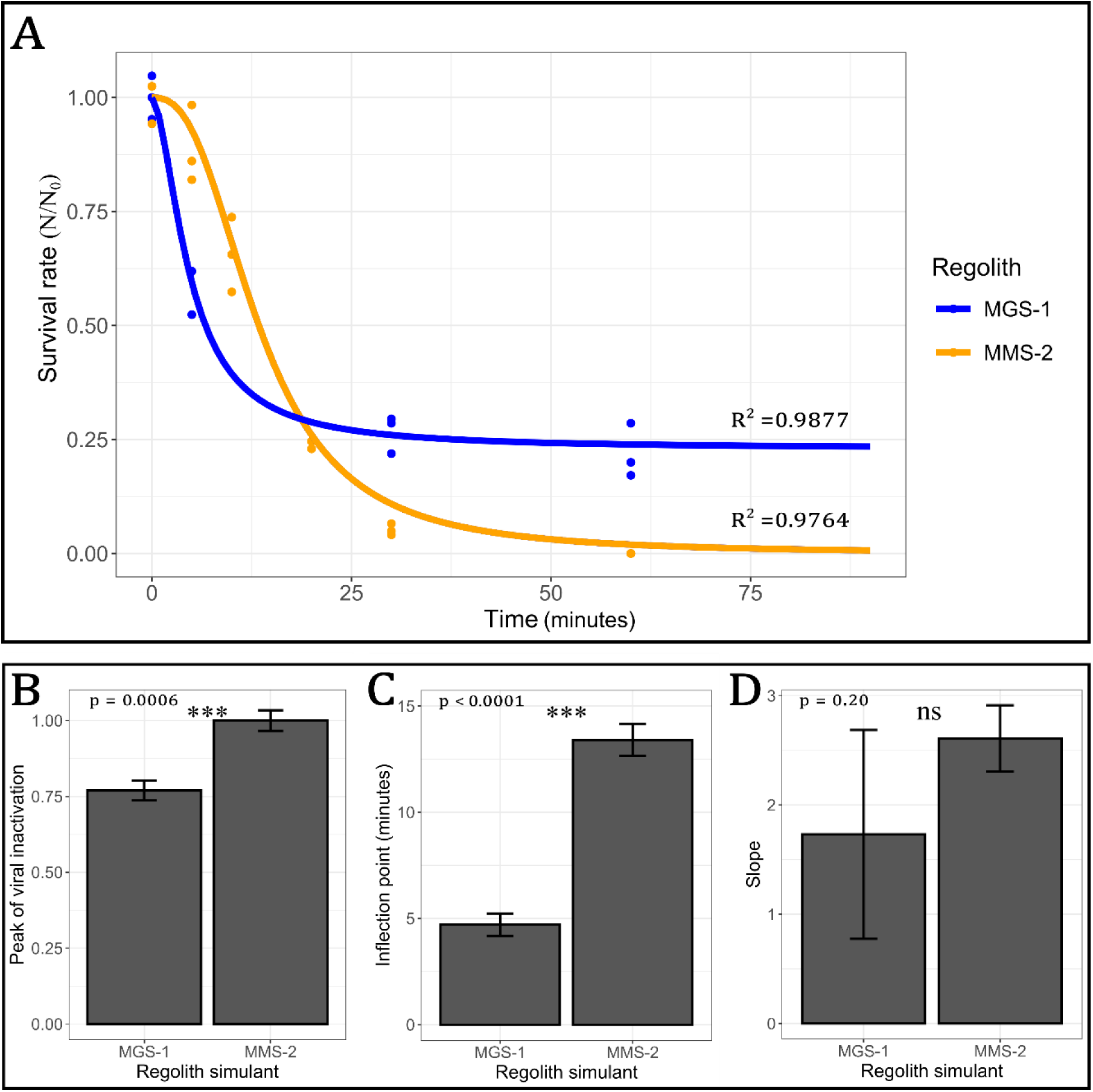
Time-dependent inactivation of bacteriophage Qβ in the presence of MGS-1 and MMS-2 at a concentration of 12.5% w/v. Incubations were performed in triplicate at 25 °C and modeled using a four-parameter logistic (4PL) function. (A) inactivation curves for both simulants. (B) maximum extent of viral inactivation. (C) inflection point (x₅₀), corresponding to the time required for 50% inactivation. (D) Slope of the curve at the inflection point. Error bars in panels B–D represent the standard deviation (SD) of each parameter as obtained through the model. Statistical significance was assessed using Welch’s *t*-test.

#### Variations in viral solution to regolith ratio

Contact with the mineral matrix is expected to be the primary driver of viral inactivation. To evaluate this effect, we tested a range of regolith-to-virus solution ratios (Figure 3). Low concentrations of both MRSs (0.5 mg per 100 µl of viral solution) resulted in minimal loss of viral infectivity. Slightly higher concentrations (2.5 mg per 100 µl) led to a modest reduction (5–10%) in viral titers. The standard concentration used throughout the study (12.5 mg per 100 µl) consistently caused 70–90% inactivation. At this concentration and time, MGS-1 led to slightly less damage than MMS-2, suggesting that the point at which both simulants lead to comparable survivals takes place before 20 min of exposure, and likely in the 15 min range. At elevated regolith concentrations (50 and 100 mg per 100 µl), viral inactivation increased dramatically, with only 1% and 0.1% of viral particles remaining, respectively. Under these conditions, the differences between the two simulants became most evident: MGS-1 consistently outperformed MMS-2, with nearly an order of magnitude higher recovery. These results suggest that MMS-2 either inactivates or retains viral particles more efficiently than MGS-1.

**Figure 3.**
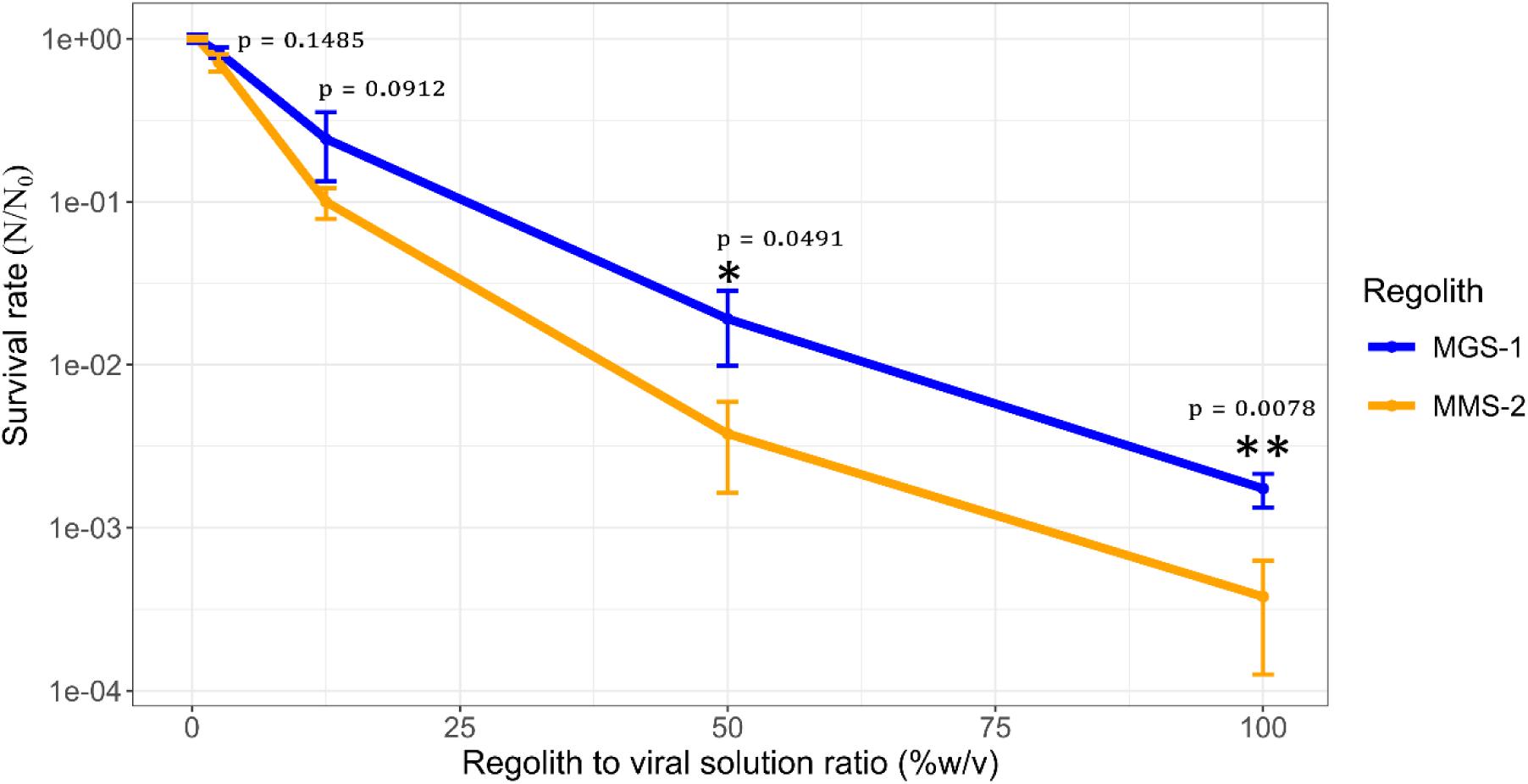
Viral infectivity of bacteriophage Qβ after 20 minutes of incubation at 25 °C with increasing concentrations of MGS-1 and MMS-2. MRS to virus solution ratios ranged from 0.5% to 100% w/v. Each point represents the mean viral titer from three independent replicates. Error bars indicate standard deviation. Statistical significance between conditions was assessed using a two-tailed two-sample Welch’s *t*-test.

#### Influence of temperature on phage-MSRs interaction

Although bacteriophage Qβ infects *E. coli*, which grows optimally at 37 °C, elevated temperatures are known to accelerate chemical reactions and may therefore enhance viral inactivation when in contact with mineral substrates. To assess this effect, viral solutions were incubated with MGS-1 and MMS-2 at four temperatures (4, 25, 30, and 37 °C) for 20 minutes and 2 hours (Figure 4). Two times were assayed to account for the different inactivation kinetics of both regoliths. Figures 4 to 7 show viral titers in barplots instead of survival rates in lineplots. This was chosen to enhance interpretability, as in most figures MMS-2 and MGS-1 have differences of several orders of magnitude, and in many cases MMS-2 led to no viral recovery.

**Figure 4.**
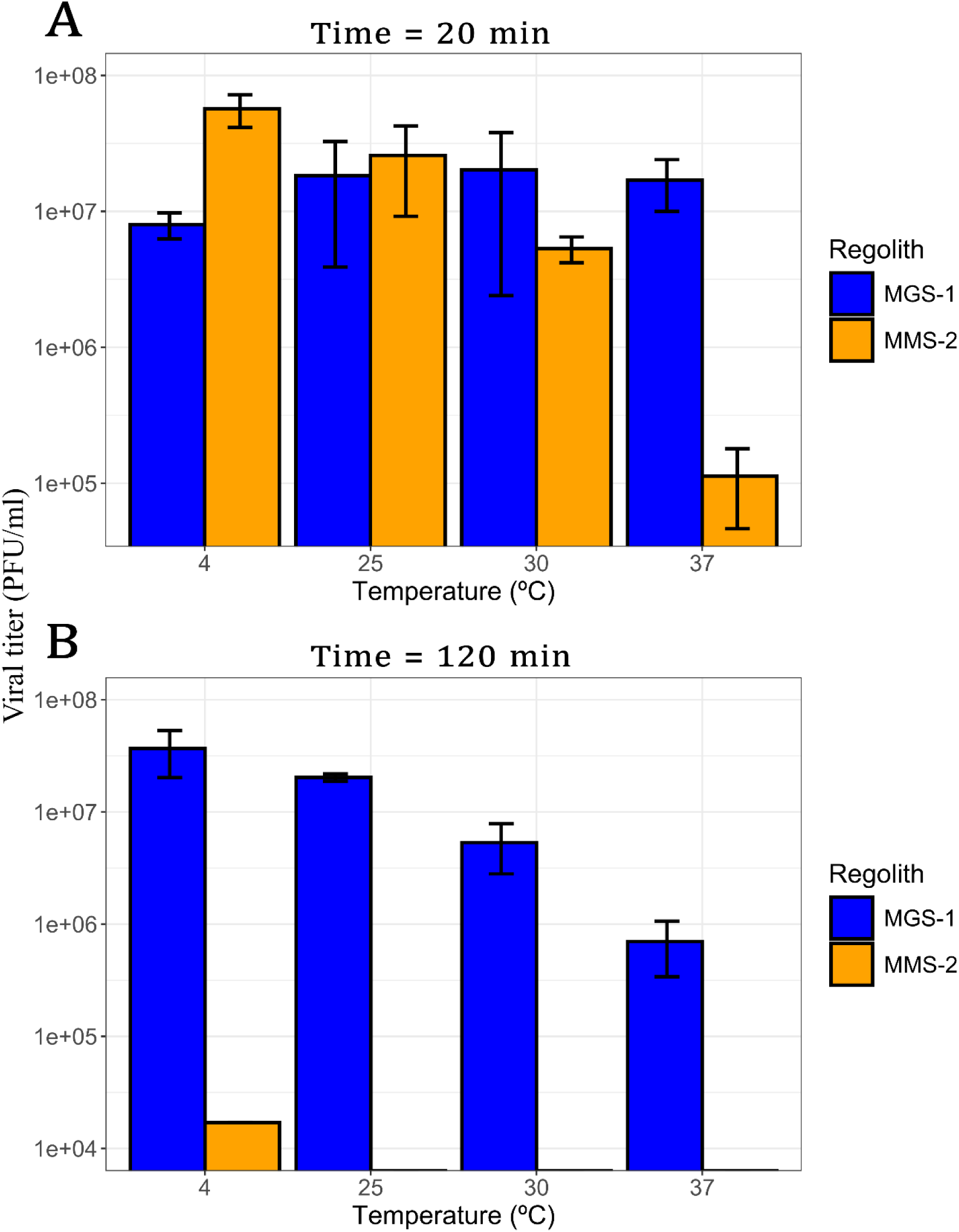
Viral infectivity of bacteriophage Qβ after incubation with MGS-1 and MMS-2, at a concentration of 12.5% w/v. (A) viral recovery after 20 minutes. (B) viral recovery after 120 minutes. In all cases the ratio of MRS to virus solution was 12.5% w/v and the initial virus titer 10^8^ PFU/ml. The temperatures assayed were 4 °C, 25 °C, 30 °C, and 37 °C. Each bar represents the mean viral titer from three independent replicates. Error bars indicate standard deviation.

After 20 minutes of exposure (Figure 4A), MGS-1 showed no significant differences in viral recovery across temperatures (*p* > 0.1), whereas MMS-2 exhibited a progressive decline in virus titers as temperature increased (*p* < 0.001). This effect was most pronounced at 37 °C, where viral survival in MMS-2 was reduced by two orders of magnitude. Following 120 minutes of exposure (Figure 4B), viral recovery from MMS-2 was negligible at all temperatures, with only minimal survival observed at 4 °C. In contrast to the short-term results, MGS-1 also showed temperature-dependent effects after prolonged exposure (*p* < 0.001), with the highest viral recovery at 4 °C and a gradual decline as temperature increased, culminating in the lowest survival at 37 °C. This pattern mirrors the short-term behavior of MMS-2, suggesting that temperature plays a critical role in modulating viral stability over time.

#### Effects of different particle sizes

Both exposed surface area and mineral composition are known to strongly influence the recovery of organic matter from mineral matrices. To further characterize this effect, bacteriophage Qβ was exposed to five different fractions of MGS-1 and MMS-2 (Figure 1) for 120 minutes at 4 °C, 25 °C, and 37 °C (Figure 5A–C). As expected from results shown in previous section, viral recovery was highest at lower temperatures for both simulants.

**Figure 5.**
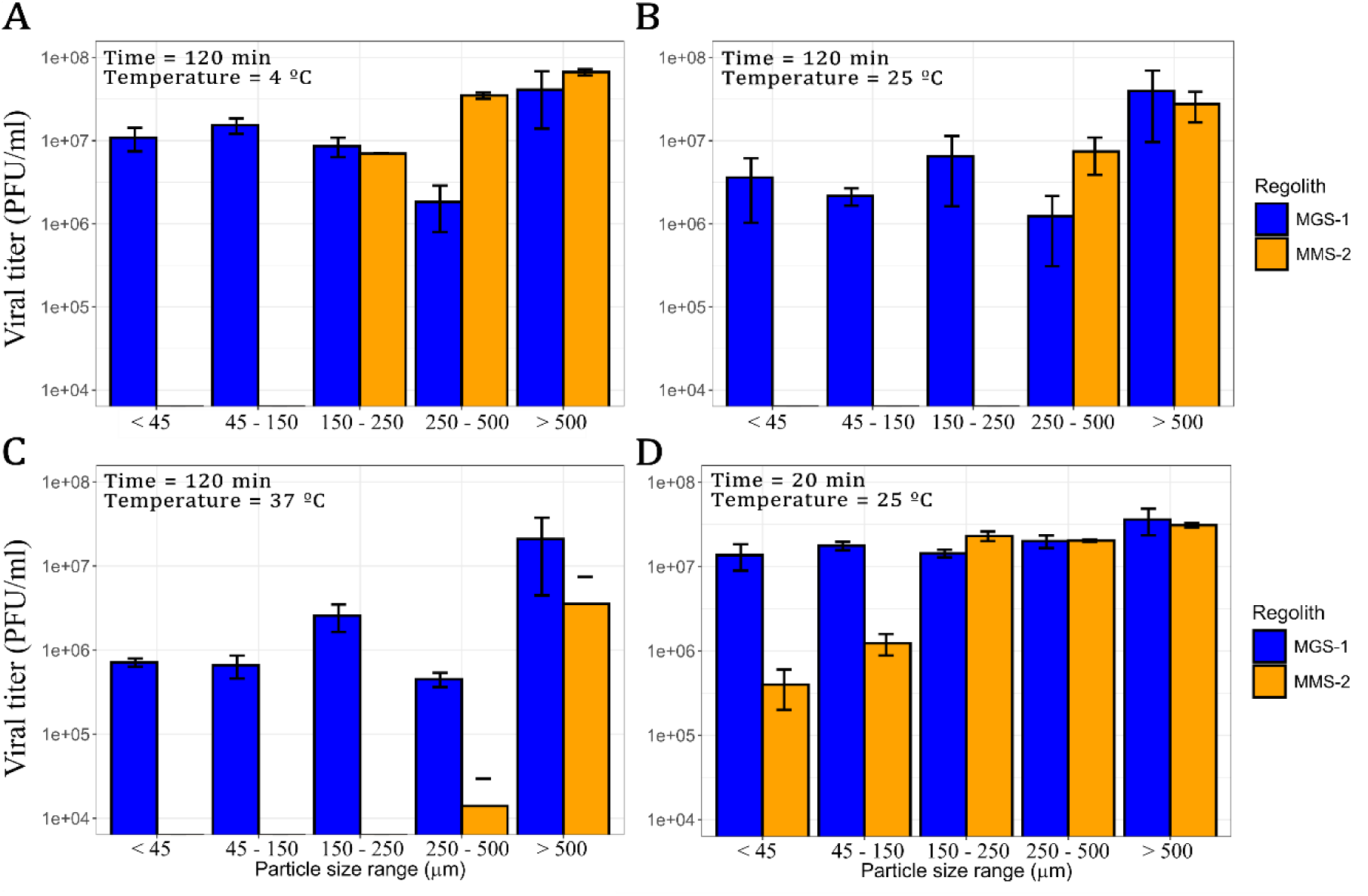
Viral infectivity of bacteriophage Qβ after incubation with five particle size fractions of MGS-1 and MMS-2 at different temperatures. In all cases the regolith concentration was 12.5% w/v and the initial viral titer 10^8^ PFU/ml. (A–C) viral recovery after 120 minutes of incubation at 4 °C, 25 °C, and 37 °C, respectively. (D) shows viral recovery after 20 minutes of incubation at 25 °C. Each point represents the mean viral titer from three independent replicates. Error bars indicate standard deviation.

In the case of MGS-1, the highest recovery was observed in particles > 0.500 mm, the lowest in the 0.250–0.500 mm range, and similar levels across the remaining fractions. Given the consistent mineral composition across MGS-1 particle sizes, viral recovery was also relatively uniform. MMS-2, however, showed a markedly different pattern. No infective viral particles were recovered from fractions < 0.150 mm. In the remaining size ranges, recovery was strongly temperature-dependent, more so than in MGS-1. At 4 °C, viral recovery in MMS-2 was comparable to MGS-1 for particles > 0.150 mm. At 25 °C, this similarity was restricted to particles > 0.250 mm, and at 37 °C, only the > 0.500 mm fraction of MMS-2 approached the recovery levels of MGS-1.

To assess short-term effects, both simulants were also incubated with the virus for 20 minutes at 25 °C (Figure 5D). Under these conditions, bulk MGS-1 and MMS-2 behaved similarly (Figure 4A), as did particles > 0.150 mm, with indistinguishable levels of viral recovery. In contrast, when particle size was < 0.150 mm, MMS-2 proved significantly more detrimental, resulting in viral titers nearly two orders of magnitude lower than those obtained with MGS-1.

### Protective role of MGS-1 against desiccation and UV radiation

Before evaluating the protective effect of Martian Regolith Simulants (MRSs) against desiccation, we first assessed the contribution of gelatin −a common stabilizing agent in viral buffers (Huang et al., 2025b)− to viral preservation. Gelatin forms films under desiccating conditions that retain moisture and protect viral particles. To avoid confounding its effects with those of the mineral substrate, we prepared two modified buffer conditions: one in which gelatin was completely removed, and another in which gelatin was replaced with bovine serum albumin (BSA), a globular protein with similar stabilizing properties. These conditions were tested both in the absence and presence of MGS-1 to evaluate their individual and combined protective effects (Figure 6).

**Figure 6.**
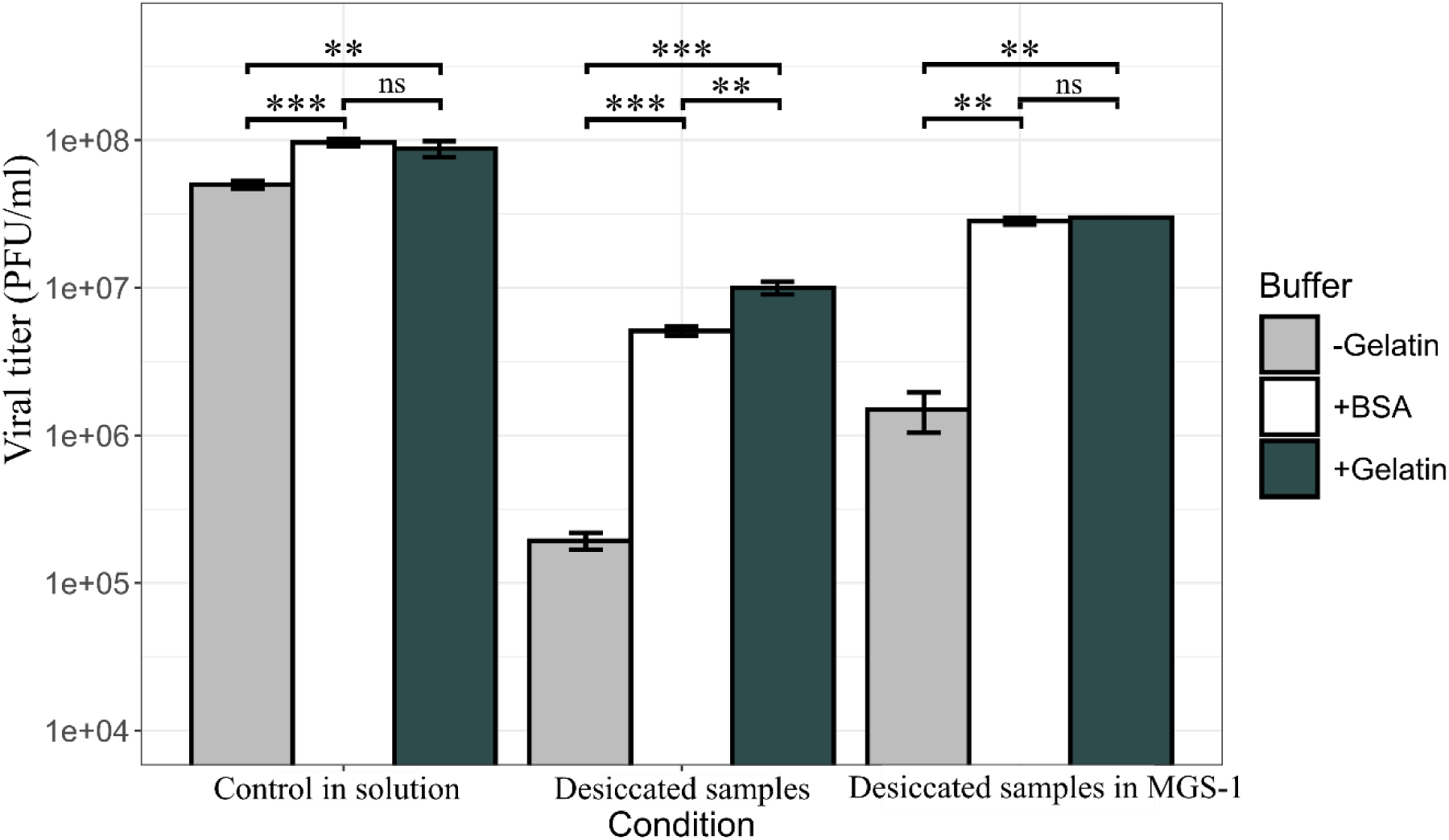
Viral infectivity of bacteriophage Qβ with and without desiccation. Desiccation was performed as described in Methods under different conditions: standard buffer containing 0.1% gelatin (black bars), standard buffer without gelatin (grey bars), and standard buffer with BSA (white bars). Each condition was tested both with and without MGS-1 to evaluate the individual and combined protective effects of proteins and mineral substrate. Each bar represents the mean viral titer from three independent replicates. Error bars indicate standard deviation. Statistical significance between conditions was assessed using Welch’s *t*-test.

**Figure 7.**
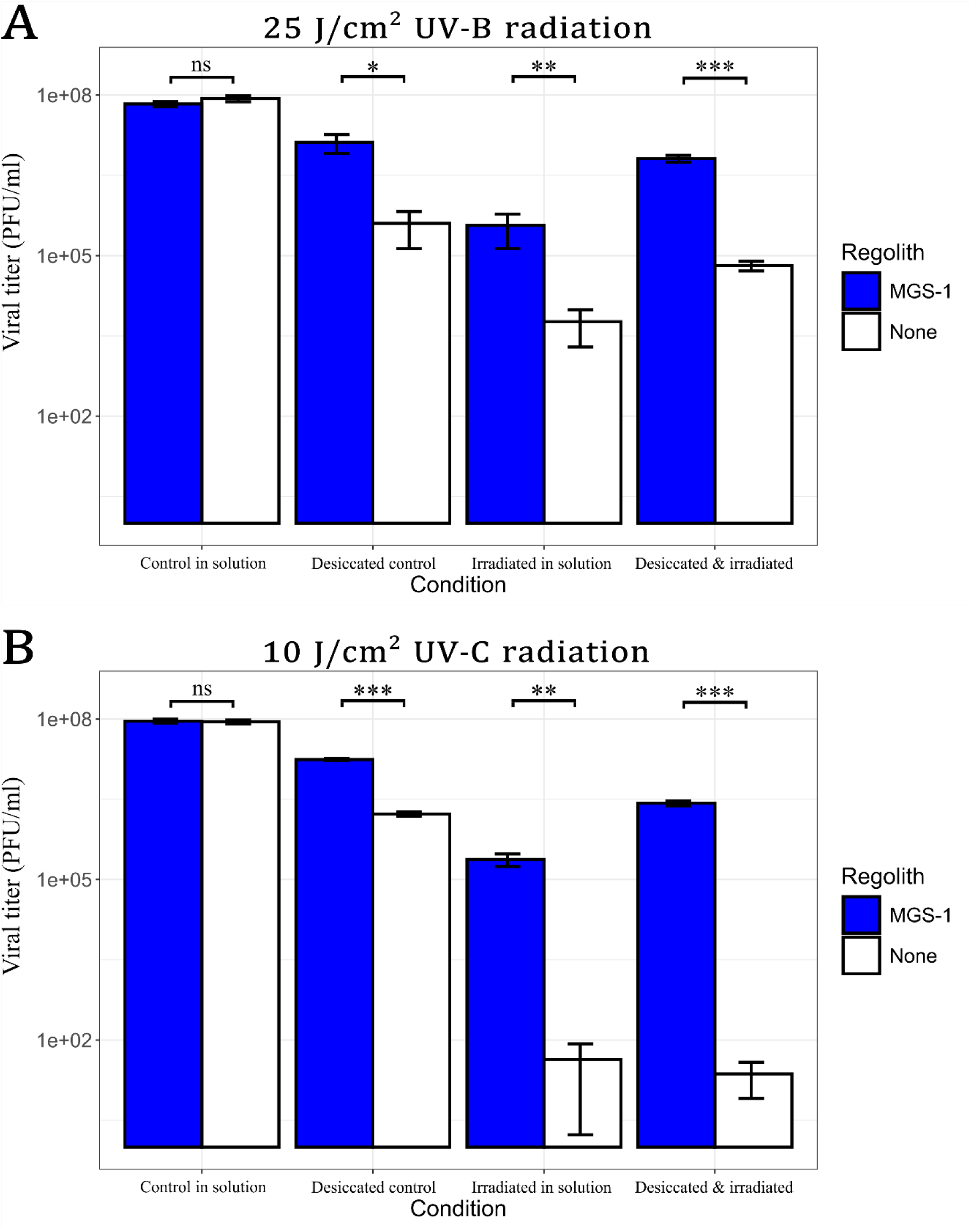
Viral infectivity of bacteriophage Qβ after exposure to ultraviolet radiation in different conditions. (A) effect of UV-B irradiation (25 J/cm²). (B) effect of UV-C irradiation (10 J/cm²). Viral samples were prepared in standard buffer and in the presence (blue bars) or the absence (white bars) of MGS-1. Exposure to radiation was performed both in solution and after desiccation. Controls were not irradiated. Each bar represents the mean viral titer from three independent replicates. Error bars indicate standard deviation. Statistical significance between conditions was assessed using Welch’s *t*-test.

Consistent with previous findings (Huang et al., 2025b), we observed that viral infectivity declined sharply in the absence of a stabilizing agent such as gelatin or BSA. Gelatin and BSA were both effective stabilizers, with gelatin leading to slightly better survivals in samples desiccated without MGS-1. Desiccation in the absence of any stabilizing agent led to drops of over one order of magnitude both with, and without MGS-1. Viruses also appeared to exhibit enhanced survival when desiccated within MGS-1. Even in samples left in solution, the absence of a stabilizing agent led to a consistent and constant decrease in viral viability. Because of this, all prior and subsequent experiments were performed using gelatin as a stabilizer. This is standard procedure and prevented artefactual viral damaging during experiments. MMS-2 was also not employed in the following experiments. This was decided as the time required to fully desiccate viruses within it was long enough to almost completely degrade them, which made further analysis and irradiation unviable.

Desiccation in the absence of MGS-1 led to almost a 3-log reduction in viral infectivity when no stabilizing protein was present. This loss was reduced to 1-log with gelatin and was only slightly higher with BSA. These results confirm the protective role of soluble proteins against desiccation-induced damage. We then evaluated the effect of MGS-1 on viral survival under desiccating conditions. In the absence of stabilizing proteins, MGS-1 reduced viral loss to <2-log. When gelatin or BSA was present, the reduction was < 1-log, indicating that MGS-1 enhances viral preservation both independently and synergistically with protein-based stabilizers. The effect of each desiccation condition was determined by comparing infectivity data with that of control samples incubated for the same duration and at the same temperature, but without undergoing desiccation.

To assess protection against UV radiation, viral samples were irradiated with UV-B and UV-C, both in the absence and presence of MGS-1 (Figure 7). Direct UV-B exposure (25 J/cm²) caused a 4-log reduction in wet samples and a 3-log reduction in desiccated samples (Figure 7A). UV-C irradiation (10 J/cm²) was more damaging, leading to a 6-log reduction in both wet and dry samples (Figure 7B) in the absence of MGS-1. Remarkably, viruses embedded within MGS-1 were significantly more resistant to UV exposure. Recovery levels were similar regardless of UV type, with only a slighter decrease in infectivity under UV-C. Desiccated viruses within MGS-1 were also better preserved than those irradiated in wet conditions, with approximately 1-log higher recovery across both UV treatments. Overall, irradiation of viruses irradiated in the while in contact with MGS-1 were better protected than those without it, regardless of UV type (UV-B and UV-C) and if the irradiation took place in dry samples or in solution.

### Protective role of MRSs against freezing and thawing cycles

The Martian surface is subject to subzero temperatures for most of the year, with transient melting of water ice occurring only under exceptional conditions (Fairén, 2010; Martínez and Renno, 2013; Sori and Bramson, 2019). Given that repeated freezing and thawing (F/T) at −20 °C is known to severely reduce the infectivity of bacteriophage Qβ (Laguna-Castro et al., 2024), we sought to investigate whether Martian Regolith Simulants could mitigate this effect.

To this end, we evaluated viral survival after successive F/T cycles in the presence of MGS-1, MMS-2, and an inert control (sea sand) (Figure 8). During the first four cycles, samples frozen in contact with MMS-2, sand, or buffer alone showed similar levels of viral loss (*p* > 0.1), whereas those incubated with MGS-1 exhibited significantly higher survival rates (cycle 1 *p* < 0.01; cycle 2 *p* < 0.001; cycle 3 *p* < 0.01; cycle 4 *p* < 0.001).

**Figure 8.**
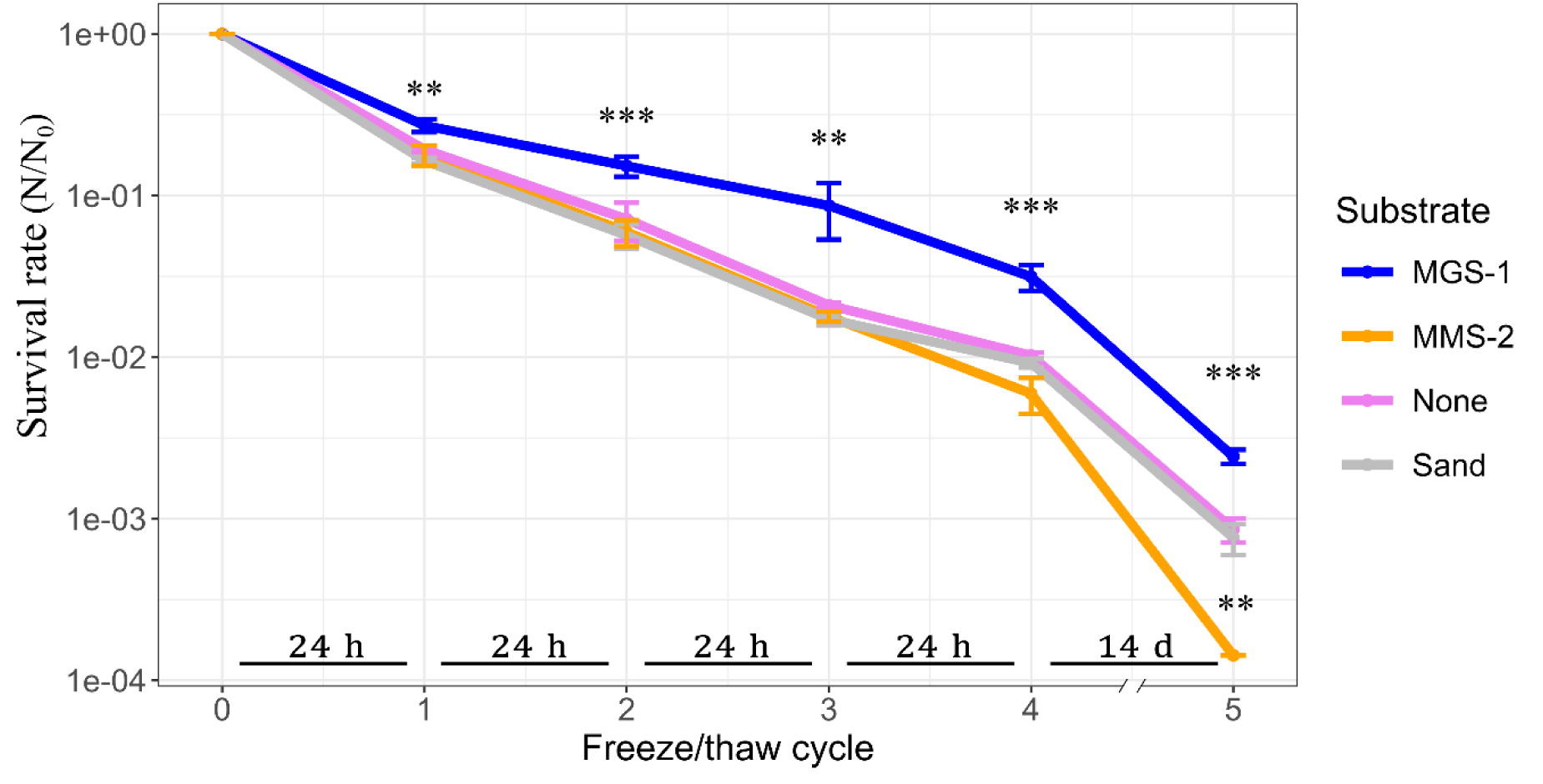
Survival rate of bacteriophage Qβ after five consecutive freeze–thaw (F/T) cycles at −20 °C. Samples were incubated with MGS-1, MMS-2, sea sand (inert control), or phage buffer alone, at a concentration of 12.5% w/v. Time spent frozen between cycles is indicated in the horizontal bars placed above the X-axis. Each point represents the mean survival rate from three independent replicates. Error bars indicate standard deviation. Statistical differences between conditions at each cycle were assessed using one-way ANOVA followed by post-hoc Dunnett’s test.

After the fifth F/T cycle −corresponding to 14 days of freezing− MGS-1 continued to outperform all other substrates (*p* < 0.001). MMS-2 was markedly more damaging than the other conditions (*p* < 0.01), and samples incubated with sand were indistinguishable from those lacking any mineral substrate (*p* > 0.1). It is important to emphasize that this final cycle takes the most duration and it is conceivable that this may be contributing to the differences becoming more pronounced. Despite these considerations these results suggest that MGS-1 provides a protective microenvironment that enhances viral robustness under repeated freeze–thaw stress.

## Discussion

Following the exposure of bacteriophage Qβ to a wide range of Mars-analog environmental stressors, clear differences emerged between the two Martian Regolith Simulants (MRSs), MGS-1 and MMS-2. In contact with MGS-1, viral particles were rapidly lost during the initial minutes (< 10 min), yet a substantial fraction remained infective even after prolonged incubation (> 120 min). This pattern is consistent with a mechanism dominated by physical adsorption, whereby viruses bind to mineral surfaces without undergoing structural degradation (Figure 2). In contrast, MMS-2 preserved viral infectivity more effectively during the early stages of exposure but led to near-complete inactivation over longer periods. This suggests that MMS-2 exerts a time-dependent virucidal effect, likely driven by chemical oxidation mediated by hematite and enhanced adsorption via gypsum. The progressive loss of viral activity in MMS-2, especially at higher regolith concentrations, points to the accumulation of reactive compounds capable of damaging viral capsids or genomes. These findings highlight the dual role of mineral substrates: while they can transiently stabilize viral particles through adsorption, their chemical reactivity may ultimately compromise viral integrity. The contrasting behaviors of MGS-1 and MMS-2 underscore the importance of mineral composition in determining the fate of viruses when in contact with mineral surfaces and suggest that even subtle differences in substrate chemistry can lead to markedly divergent outcomes.

Adsorption, as a physical process, is generally not strongly influenced by temperature. While elevated temperatures increase the kinetic energy of viral particles—enhancing their collision frequency with mineral surfaces—they can also promote desorption through increased molecular motion and potential destabilization of the viral capsid. In contrast, lower temperatures tend to favor stable adsorption, as previously reported (Syngouna and Chrysikopoulos, 2010). Chemical processes such as oxidation, however, are highly temperature-dependent, with reaction rates accelerating as temperature rises. The pronounced temperature sensitivity observed in MMS-2 again suggests that chemical oxidation −likely mediated by hematite− is the dominant mechanism driving viral inactivation, even over short exposure periods (20 min) (Figure 4A). In contrast, MGS-1 showed minimal variation in viral survival across temperatures during short incubations, reinforcing the idea that physical adsorption, rather than chemical degradation, governs its interaction with viral particles (Figure 4A). Over extended timeframes (120 min) (Figure 4B), viral inactivation in MMS-2 became nearly complete, regardless of temperature, indicating that oxidative damage accumulates over time. Interestingly, MGS-1 began to show temperature-dependent effects under these longer exposures, suggesting that although primarily inert, some minor chemical interactions may contribute to viral degradation in this substrate. These findings underscore the importance of both mineral composition and exposure duration in shaping viral fate under Mars-analog conditions.

The Martian regolith, however, is composed of particles of a wide range of sizes and topologies. Different particle sizes were expected to and exhibited distinctive damage profiles when viruses were exposed to them (Figure 5). Over short (20 min) periods of time (Figure 5D), MGS-1 led to little damage irrespective of particle size, while the smaller particles of MMS-2 heavily damaged viruses in contact with them. Gypsum and hematite were abundant in these smaller particles of MMS-2 and likely drove viral loss through fast adsorption to gypsum and oxidation by hematite. Over longer periods of time (Figure 5A-C), viruses within MGS-1 show decreased survival. This is, however, mostly independent of particle size. In the case of MMS-2, no viruses could be recovered from the two smallest, gypsum-rich particle sizes after 2 hours. As temperature increased, viral recovery decreased uniformly in MGS-1 (except for the > 500 µm range). In MMS-2, however, an increase in temperature led to changes in the threshold at which practically no viruses could be recovered, it being the 45 to 150 µm range at 4 °C, the 150 to 250 µm range at 25 °C, and the 250 to 500 µm range at 37 °C. Overall, viral recovery appeared greatly dependent on particle size in MMS-2, where composition is more varied and the increased availability of exposed surface area and oxidative minerals appeared to drive viral inactivation at lower particle size ranges. This was not the case in MGS-1, whose consistent composition and relative lack of oxidative components leads to a tamer chemical environment.

The presence of soluble proteins as stabilizing agents proved essential for protecting bacteriophage Qβ from desiccation and freeze-induced damage, increasing viral survival by up to two orders of magnitude. Although both gelatin and BSA were effective, gelatin consistently offered slightly superior protection. This suggests that the protective effect is not strictly dependent on the specific protein used, but rather on the general capacity of globular proteins to form hydration shells or structural matrices that buffer viral particles against environmental stress. It is important to note that viruses rarely exist in isolation in natural environments. Instead, they are embedded within complex biological matrices that include salts, proteins, lipids, and cellular debris from host organisms. In this context, the high concentrations of salts and proteins present in the culture medium may contribute to viral stabilization (Huang et al., 2025), while residual cellular debris from the infection process may further enhance protection by physically shielding viral particles or facilitating aggregation (Laguna-Castro et al., 2023). These findings underscore the relevance of simulating realistic biological conditions when assessing viral survivability, particularly in astrobiological scenarios where viruses may be embedded in organic or mineral-rich microenvironments.

Additionally, desiccation in the presence of MGS-1 consistently led to higher viral survival across all tested conditions. This observation aligns with previous findings by Zaccaria et al. (Zaccaria et al., 2024), who reported that desiccation-sensitive microorganisms were better able to withstand dehydration when embedded in Martian regolith simulants, and in some cases only survived desiccation when such substrates were present. One plausible explanation is the hygroscopic nature of the regolith particles, which may retain residual moisture and create microenvironments that buffer biological entities against extreme dryness (Jänchen et al., 2014). The water-retention capacity of Martian regolith simulants has broader implications beyond viral preservation. It reinforces the hypothesis that certain mineral substrates on Mars could have supported transient habitable niches in the planet’s past, particularly during epochs when liquid water was more abundant (Mansilla et al., 2023). Moreover, this property is highly relevant for the design of future sampling strategies in Mars missions, as it suggests that regolith particles may act as reservoirs of biological material or biomarkers (Zorzano et al., 2024). Understanding how desiccation interacts with mineral composition is therefore crucial not only for astrobiology, but also for planetary protection and the interpretation of potential biosignatures.

Irradiation of viral particles in solution consistently resulted in lower survival rates compared to those embedded in dried substrates. This disparity may be explained by the dynamic behavior of viruses in liquid media, where recirculation and diffusion can bring particles repeatedly to the surface, increasing their exposure to damaging radiation. In contrast, desiccated samples likely restrict viral mobility, reducing the probability of direct irradiation. Moreover, the presence of water in liquid samples facilitates the generation of reactive oxygen species (ROS), particularly in the presence of iron ions such as those found in hematite. These ROS—including hydroxyl radicals and hydrogen peroxide—are highly reactive and capable of inflicting severe damage to nucleic acids and capsid proteins (He et al., 2016; Kaur et al., 2016). The synergistic effect of water and iron-rich minerals in promoting oxidative stress may therefore explain the pronounced loss of viral infectivity observed in wet conditions. These findings highlight the importance of both hydration state and mineral composition in modulating the impact of radiation on viral particles under Mars-analog conditions.

MGS-1 is currently regarded as the “gold standard” among Martian regolith simulants (Cannon et al., 2019; Nababan et al., 2025; Sun et al., 2023), due to its compositional stability, reproducibility, and extensive experimental validation. It has been widely employed in studies ranging from biomarker preservation to plant cultivation (Arribas Tiemblo et al., 2025; Macário et al., 2022). In our experiments, MGS-1 consistently emerged as the least damaging substrate, exhibiting low oxidative potential, strong adsorptive and hygroscopic properties, and notable protective effects against UV radiation. However, its lack of iron oxides may render it less representative of the more reactive conditions found on the Martian surface. In contrast, MMS-2 contains substantial amounts of iron oxides—primarily hematite—and may better reflect the composition of specific Martian terrains. This compositional difference, however, translated into significantly reduced viral preservation and protection. Our findings suggest that even slight variations in the mineralogical makeup of Martian regolith, particularly in iron oxide content and adsorptive minerals, can lead to pronounced differences in viral survival. Overall, our findings reveal marked and heterogeneous responses of bacteriophage Qβ to different regolith simulants, underscoring the need to identify—or engineer through tailored mixtures—new, more representative natural simulants (Costa et al., 2024).

Early experiments exposing viruses to space conditions date back to the Gemini IX and XII missions, where bacteriophage T1 and tobacco mosaic virus (TMV) were subjected to the full space environment (Horneck et al., 2010). These studies revealed the strong inactivating effect of space exposure (Hotchin et al., 1968), and notably, the survival of T1 improved when samples were shielded with a thin layer of aluminum. This suggested that non-penetrating radiation—likely solar UV or soft X-rays—was primarily responsible for viral inactivation. In our study, we observed a similar protective effect when samples were covered with Martian regolith simulant MGS-1. Regardless of the quantity used, MGS-1 provided significant shielding against UV-B and UV-C radiation, reinforcing its potential role as a natural protective matrix. These results not only align with historical findings but also highlight the relevance of regolith composition in modulating the impact of extraterrestrial radiation on biological entities.

In the broader context of astrobiology, our results suggest that the mineralogical diversity of planetary surfaces may play a decisive role in shaping the fate of viral particles, whether as relics of past biological activity or as active entities. The differential survival of bacteriophage Qβ across simulants underscores the importance of substrate composition in modulating viral stability and invites further exploration into how such interactions might unfold on Mars, icy moons, or other bodies of interest.

## Acknowledgements

The authors would like to thank Maite Fernández-Sampedro for her assistance in the performance of X-Ray Diffraction (XRD) analysis. We also thank Dr. Eduardo González Pastor and his team at Centro de Astrobiología (CAB) for their input on the UV-C irradiation of samples.

## Authorship Contribution Statement

**Miguel Arribas Tiemblo:** Conceptualization, Formal analysis, Investigation, Methodology, Validation, Visualization, Writing – original draft.

**Alicia Rodríguez-Moreno:** Conceptualization, Formal analysis, Investigation, Methodology, Validation, Writing – original draft.

**Felipe Gómez:** Funding acquisition, Methodology, Validation, Supervision, Writing – review and editing.

**Ester Lázaro:** Conceptualization, Formal analysis, Funding acquisition, Methodology, Validation, Supervision, Writing – review and editing.

